# Extracellular vesicle-derived miRNA-mediated cell-cell communication inference for single-cell transcriptomic data with miRTalk

**DOI:** 10.1101/2024.07.07.602386

**Authors:** Xin Shao, Chengyu Li, Jingyang Qian, Haihong Yang, Xinyu Yang, Jie Liao, Xiao Xu, Xiaohui Fan

## Abstract

MicroRNAs are released from cells in extracellular vesicles (EVs), representing an essential mode of cell-cell communication (CCC) via an inhibitory effect on gene expression. The advent of single-cell RNA-sequencing (scRNA-seq) technologies has ushered in an era of elucidating EV-derived miRNA-mediated CCC. However, the lack of computational methods to infer such CCC poses an outstanding challenge. Herein, we present miRTalk (https://github.com/multitalk/miRTalk), a pioneering framework for inferring EV-derived miRNA-mediated CCC with a probabilistic model and a curated database, miRTalkDB, which includes EV-derived miRNA-target associations. The benchmarking against simulated and real-world datasets demonstrated the remarkable accuracy and robustness of miRTalk. Subsequently, we employed miRTalk to uncover the in-depth CCC mechanisms underlying three disease scenarios. In summary, miRTalk represents the first approach for inferring EV-derived miRNA-mediated CCC with scRNA-seq data, providing invaluable insights into the CCC dynamics underpinning biological processes.

## Background

MicroRNAs (miRNAs) comprise a class of small RNA molecules that predominantly orchestrate numerous cellular processes through post-transcriptional gene silencing in eukaryotes[1]. Over the past decade, miRNAs have been discovered within extracellular environments and have demonstrated their capacity to facilitate functional cell-cell communication (CCC)[2]. MiRNAs produced and released from one cell via extracellular vesicles (EVs) such as exosomes and microvesicles are captured by distant cells, subsequently leading to alterations in gene expression, cellular behavior, and function[2]. To date, EV-derived miRNAs have achieved widespread recognition as a distinctive mode of CCC, with mounting evidence demonstrating the pivotal role of EV-derived miRNA-mediated CCC in the emergence and progression of diseases[3, 4]. For instance, Wang et al. discovered that the depletion of miR-9-3p in adipose tissue-derived EVs significantly mitigates their deleterious effects on cognitive function in high-fat diet mice or patients with diabetes, thus unveiling the crucial role of the adipose tissue-neuron communication mediated by EV-derived miR-9-3p in promoting synaptic loss and cognitive impairment[5]. He et al. reported that EV-derived miR-223 from neutrophils functioned to impede hepatic inflammatory and fibrogenic gene expression, revealing the neutrophil-to-hepatocyte communication via miR-223-enriched EV transfer in the alleviation of nonalcoholic steatohepatitis[6]. Nonetheless, due to the limitations of related technologies and the pervasive cellular heterogeneity within a given tissue, the large-scale systematic investigation of EV-derived miRNA-mediated CCC among heterogeneous cell populations becomes substantially difficult, severely constraining the advancement of this field.

Fortunately, increasing progress in high-throughput single-cell RNA-sequencing (scRNA-seq) technologies has facilitated the classification of thousands of cells in a single assay based on transcriptome profiling of coding and non-coding genes[7, 8], enabling the elucidation of EV-derived miRNA-mediated CCC mechanisms underlying physiological and pathological processes at single-cell resolution[9, 10]. As an illustration, Ramanujam et al. identified the cardiac macrophage-derived miR-21 as vital in the transition from quiescent fibroblasts to myofibroblasts through single-cell analysis of pressure-overloaded hearts from mice featuring macrophage-specific ablation of miR-21, thereby revealing the involvement of miR-21 in guiding macrophage-fibroblast communication regarding myocardial homeostasis and malady-associated remodeling[9]. Zhang et al. performed scRNA-seq analyses of human intrahepatic cholangiocarcinoma and neighboring samples and found that cholangiocarcinoma cell-derived exosomal miR-9-5p engendered elevated expression of IL-6 in vascular cancer-associated fibroblasts (vCAFs) to expedite tumor progression, underscoring the significance of miR-9-5p-dependent CCC between cholangiocarcinoma cells and vCAFs within the tumor microenvironment (TME)[10]. Nevertheless, the absence of a computational approach for inferring EV-derived miRNA-mediated CCC based on scRNA-seq data presents a great challenge.

In this study, we developed a distinctive computational method, namely miRTalk (https://github.com/multitalk/miRTalk), with a bespoke probabilistic model to infer EV-derived miRNA-mediated CCC based on scRNA-seq data. By integrating experimentally verified data of EV-derived miRNAs and their corresponding miRNA-target interactions from diverse repositories, we have curated a comprehensive database called miRTalkDB that encompasses EV-derived miRNA-target associations for Homo sapiens, Mus musculus, and Rattus norvegicus. With miRTalkDB as the underlying reference, the probabilistic model aims to predict the likelihood of EV-derived miRNA-mediated CCC based on the elevation of miRNA expression in sender cells and the reduction of cognate miRNA and target gene expression in receiver cells. The performance of miRTalk was validated through simulated and real-world benchmarking datasets. Furthermore, in exploring scRNA-seq data from three disease scenarios— human glioblastoma, mouse kidney fibrosis, and rat fatty liver transplantation— miRTalk elucidated the intricate, yet hitherto unseen, EV-derived miRNA-mediated CCC. In summary, as the first of its kind, miRTalk paves the way for the computational assessment of EV-derived cellular crosstalk from scRNA-seq data, offering unprecedented insights into the CCC dynamics that underlie biological processes.

## Results

### Overview of the miRTalk method

First, we curated an EV-derived miRNA-target database named miRTalkDB as the fundamental reference. As depicted in **Fig. 1a**, we collected and manually verified the EV-derived miRNAs of Homo sapiens, murine, and Rattus norvegicus miRNAs from exiting databases—ExoCarta[11], Vesiclepedia[12], EVmiRNA[13]—and primary litertures, thereby ensuring the inclusion of miRNAs present within these established repositories that record miRNAs found in diverse extracellular vesicle categories. By incorporating experimentally verified information on miRNA-target interactions derived from miRTarBase[14], a curated EV-derived miRNA-target database was constructed. Specifically, miRTalkDB features 2,567; 935; and 179 EV-derived miRNAs, and 495,217; 48,480; and 632 miRNA-target interactions for Homo sapiens, Mus musculus, and Rattus norvegicus, respectively (**Fig. 1b**).

**Fig. 1.**
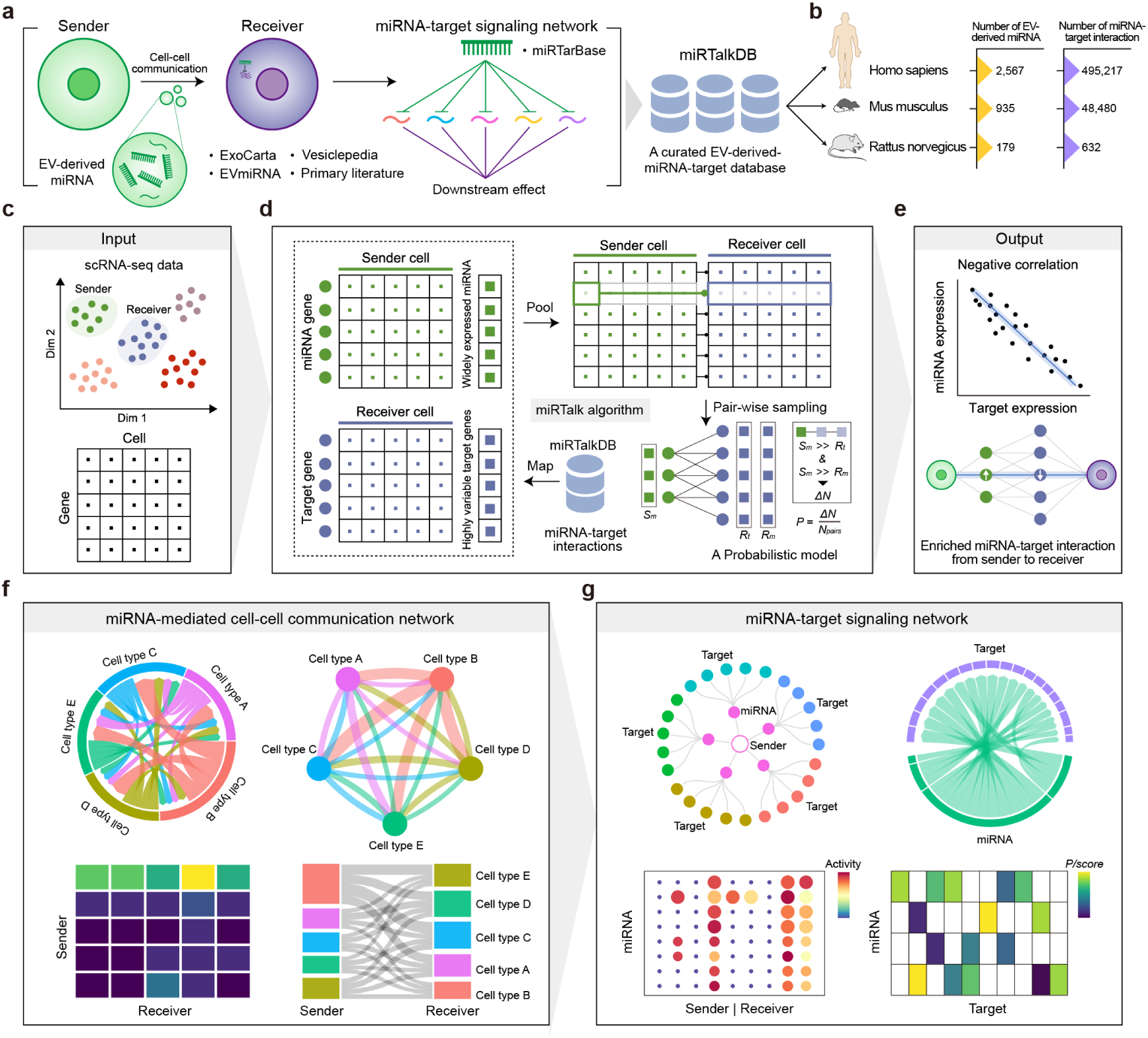
Schematic representation of the miRTalk workflow and visualization. **(a)** Construction of the miRTalkDB based on the EV-derived miRNAs catalogued in ExoCarta, Vesiclepedia, EVmiRNA, and primary literatures. By integrating experimentally verified data on miRNA-target interactions from miRTarBase, a curated EV-derived miRNA-target database, namely miRTalkDB, was constructed as the underlying reference for the subsequent inference. **(b)** Statistics of the miRTalkDB including the number of EV-derived miRNAs and miRNA-target interactions for Homo sapiens, Mus musculus, and Rattus norvegicus. **(c)** Input data of miRTalk including a scRNA-seq data matrix and corresponding cell type annotation for each cell. **(d)** Detailed diagram of the miRTalk algorithmic process: for a given sender-receiver pair, the widely expressed miRNAs and highly variable target genes were identified based on expression levels in both cell types. Based on pairwise sampling, a probabilistic model was incorporated, assuming CCC occurs when the sender cells’ miRNA demonstrates high expression, while coinciding with low expression of the same miRNA and corresponding target genes in receiver cells. **(e)** Output data of miRTalk encompassing an enrichment of EV-derived miRNAs and associated miRNA-target interactions between sender and receiver cells. **(f)** Examination of miRNA-mediated CCC networks using chord, circle, Sankey, and heatmap visualizations, which illustrate the quantity and score of miRNA-target interactions from sender cells to receiver cells. **(g)** Visualizations of miRNA-target signaling network from sender cell types to receiver cell types, represented by chord, circle, bubble, and heatmap plots.

Next, we designed a probabilistic model to infer EV-derived miRNA-mediated CCC based on scRNA-seq data (**Fig. 1c**). This model presupposes that CCC occurs when miRNA expression levels are heightened in sender cells, concomitant with low expression of the same miRNA and corresponding target genes in receiver cells. In practice, we filtered widely expressed miRNAs and highly variable target genes in sender and receiver cells, by refering to the miRNAs and their corresponding target genes in miRTalkDB. Then, we performed sender-receiver pair sampling and identified pairs exhibiting elevated miRNA expression in sender cells relative to receiver cells, alongside downregulated target genes in receiver cells compared to other cells— thereby calculating the probability of miRNA-target interactions, detailed in the scetion of Methods. This approach ensures that inferred target genes in receiver cells demonstrate the inhibitory influence of EV-derived miRNA originating from sender cells. Consequently, we enriched significant miRNA-target interactions mediating CCC from sender to receiver cells (**Fig. 1d**).

Since certain mature miRNAs are encoded by multiple miRNA genes, we aggregated miRNA activity and miRNA-target interaction scores and selected the maximum probability of the miRNA-target interaction for the evaluation of EV-derived miRNA-mediated CCC and miRNA-target signaling networks (**Fig. 1f-g**). Through the integration of chord, circle, bubble, and heatmap plots, miRTalk enables the visualization of CCC mediated by EV-derived miRNA-target interactions from multiple perspectives.

### Robust performance of miRTalk on simulated benchmarking data

The performance of miRTalk was evaluated utilizing simulated benchmark datasets through the ESCO[15] R package for the generation of simulated scRNA-seq data with several co-expressed genes. In light of the negative regulation of miRNAs on their target genes, we reasoned that miRNA-target interactions distinguished by highly expressed miRNA genes in sender cells and lowly expressed target genes in receiver cells are true predictions, whilst interactions characterized by lowly expressed miRNA genes in sender cells and highly expressed target genes in receiver cells were deemed false ones (**Fig. 2a**). In practice, we simulated 1,000 cells encompassing 510 sender cells and 490 receiver cells with 500 miRNA genes matched to 500 target genes, and denoted the positive and negative samples from the inversely correlated miRNA-target interactions conforming to the ground truth as described above (**Fig. 2b**). Consequently, miRTalk accurately predicted the most positive and negative samples with AUROC and AUPRC reaching 0.93 and 0.98, respectively (**Fig. 2c**). By modulating key parameters such as gene number (1,000, 800, 600, 400, and 200), ligands/receptors coverage (50%, 40%, 30%, 20%, and 10%), cell number (1,000, 800, 600, 400, and 200), and sender/receiver ratio (8:2, 6:4, 5:5, 4:6, and 2:8), we constructed 20 different simulated datasets under four discrete contexts to systematically evaluate the performance of miRTalk. Accordingly, miRTalk achieved a high AUROC and AUPRC on each simulated dataset (**Fig. 2d**), showcasing robust performance in simulated datasets with the different gene number, ligands/receptors coverage, cell number, and sender/receiver ratio.

**Fig. 2.**
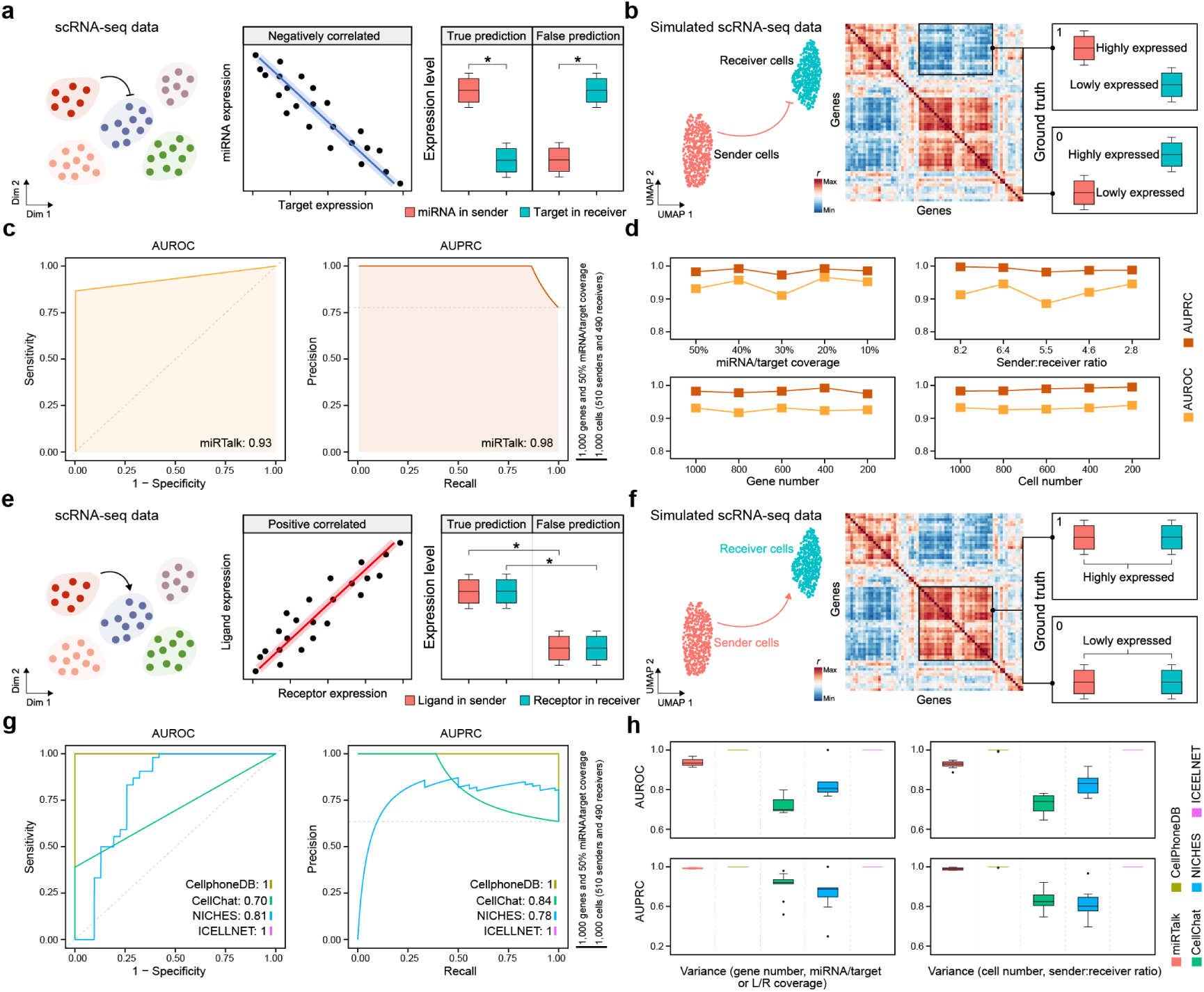
Robust performance of miRTalk on simulated benchmarking data. **(a)** Hypothesis regarding the true and false predictions of miRNA-target interactions governing the sender-receiver communications. **(b)** Simulating a scRNA-seq dataset that comprises communicative sender and receiver cell types. Among the negatively correlated pairs, the ground truth maintains that the miRNA-target interactions with highly expressed miRNA genes in sender cells and lowly expressed target genes in receiver cells are positive samples, while interactions with lowly expressed miRNA genes in sender cells and highly expressed target genes in receiver cells are negative samples. **(c)** The performance of miRTalk on the simulated data with 510 sender and 490 receiver cells and 50% miRNA/target coverage among 1000 genes, evaluated with AUROC and AUPRC. **(d)** Benchmarking analysis of miRTalk on 20 different simulated datasets under four distinct contexts through the variation of key parameters such as gene number (1,000, 800, 600, 400, and 200), ligands/receptors coverage (50%, 40%, 30%, 20%, and 10%), cell number (1,000, 800, 600, 400, and 200), and sender/receiver ratio (8:2, 6:4, 5:5, 4:6, and 2:8). **(e)** Hypothesis about the true and false predictions of LR interactions mediating the sender-receiver communications. **(f)** Simulating a scRNA-seq dataset containing communicative sender and receiver cell types. Among the positively correlated pairs, the ground truth holds that the LR interactions with highly expressed ligand and receptor genes respectively in sender and receiver cells are positive samples, while interactions with lowly expressed ligand and receptor genes respectively in sender and receiver cells are negative samples. **(g)** The performance of CellPhoneDB, CellChat, NICHES, and ICELLNET on the simulated data with 510 sender and 490 receiver cells and 50% L/R coverage among 1000 genes, evaluated with AUROC and AUPRC. **(h)** Benchmarking analysis of miRTalk, CellPhoneDB, CellChat, NICHES, and ICELLNET on 20 different simulated datasets as described above.

It is noted that the LR-based cell-cell communication inference rests on the prevalent hypothesis pertaining to the elevated expression of positively correlated ligand and receptor genes. Hence, we reasoned that LR interactions with highly expressed ligand and receptor genes respectively in sender and receiver cells represented positive samples, while interactions with lowly expressed ligand and receptor genes respectively in sender and receiver cells were considered as negative samples (**Fig. 2e**). In practice, we applied the same simulated datasets but replaced miRNA and target genes with virtual ligand and receptor genes, respectively, to maintain consistency and comparability with the simulation for evaluating miRTalk and other methods, and labeled the positive and negative samples from the positively correlated LR interactions according to the ground truth as described above (**Fig. 2f**). Notably, the established extant LR-based CCC inference methods, i.e., CellPhoneDB[16], CellChat[17], NICHES[18], and ICELLNET[19], all accurately predicted the most positive and negative samples with AUROC reaching 1, 0.70, 0.81, and 1, respectively, and AUPRC achieving 1, 0.84, 0.78, and 1, respectively (**Fig. 2g**). The accurate prediction achieved by these methods is a testimony to the reliability of the simulated process and the corresponding datasets for assessing the performance of inference methods of CCC mediated by either LR or miRNA-target interactions. Moreover, leveraging the 20 different simulated datasets under four distinct contexts as elucidated above, we compared the performance of miRTalk and other methods in predicting the positive and negative samples. Remarkably, miRTalk showed comparable AUROC and AUPRC concerning the gene and cell variance found in the simulations (**Fig. 2h**), demonstrating the impressive performance of miRTalk in inferring the EV-derived miRNA-mediated CCC.

### Prediction of miRTalk consistent with the literature evidence on real datasets

Given the well-established role of oncogenic miRNAs and tumor suppressor (TS)-miRNAs within the TME, it is anticipated that in the absence of external intervention, the expression of oncogenic miRNAs will supersede that of TS-miRNAs within malignant cells (**Fig. 3a**). Accordingly, we collected scRNA-seq datasets of 12 neoplastic specimens, derived from bladder[20], cholangiocarcinoma[10], and ovarian[21] malignancies, in order to infer the relative activity of miRNAs within cancerous cells (**Supplementary Fig. S1a-c**). Specifically, miR-24-3p is known to propagate tumorigenesis[22–24], while miR-29a-3p has been implicated in the suppression of malignant neoplasms across the aforementioned cancer types[25–27]. Remarkably, our analysis revealed a significant upsurge in miR-24-3p activity relative to miR-29a-3p within malignant cells across all 12 tumor specimens (**Fig. 3b, Supplementary Fig. S1d**), mirroring findings described within the literature. In bladder cancer specifically, we observed markedly increased activities of oncogenic miR-24-3p and miR-940 compared to the well-documented tumor suppressor miRNAs miR-29a-3p, miR-142-3p, miR-142-5p, and miR-335-5p (**Fig. 3c, Supplementary Fig. S1e**), which aligns with their respective roles in the tumor pathogenesis and progression[28–31]. Within the cholangiocarcinoma, the TS-miRNAs miR-142-5p and miR-29a-3p consistently exhibited diminished activities compared to oncogenic miR-24a-3p within cholangiocarcinoma cells[23, 26, 32] (**Supplementary Fig. S1f**). Though variabilities existed between the six ovarian cancer patient samples, the overall heightened activity of oncogenic miR-24a-3p was evident in comparison to well-established TS-miRNAs – namely, miR-29a-3p, miR-34a-5p, miR-10a-5p, and miR-92b-3p – reflecting trends supported by existing literature[24, 27, 33–35] (**Supplementary Fig. S1g-h**).

**Fig. 3.**
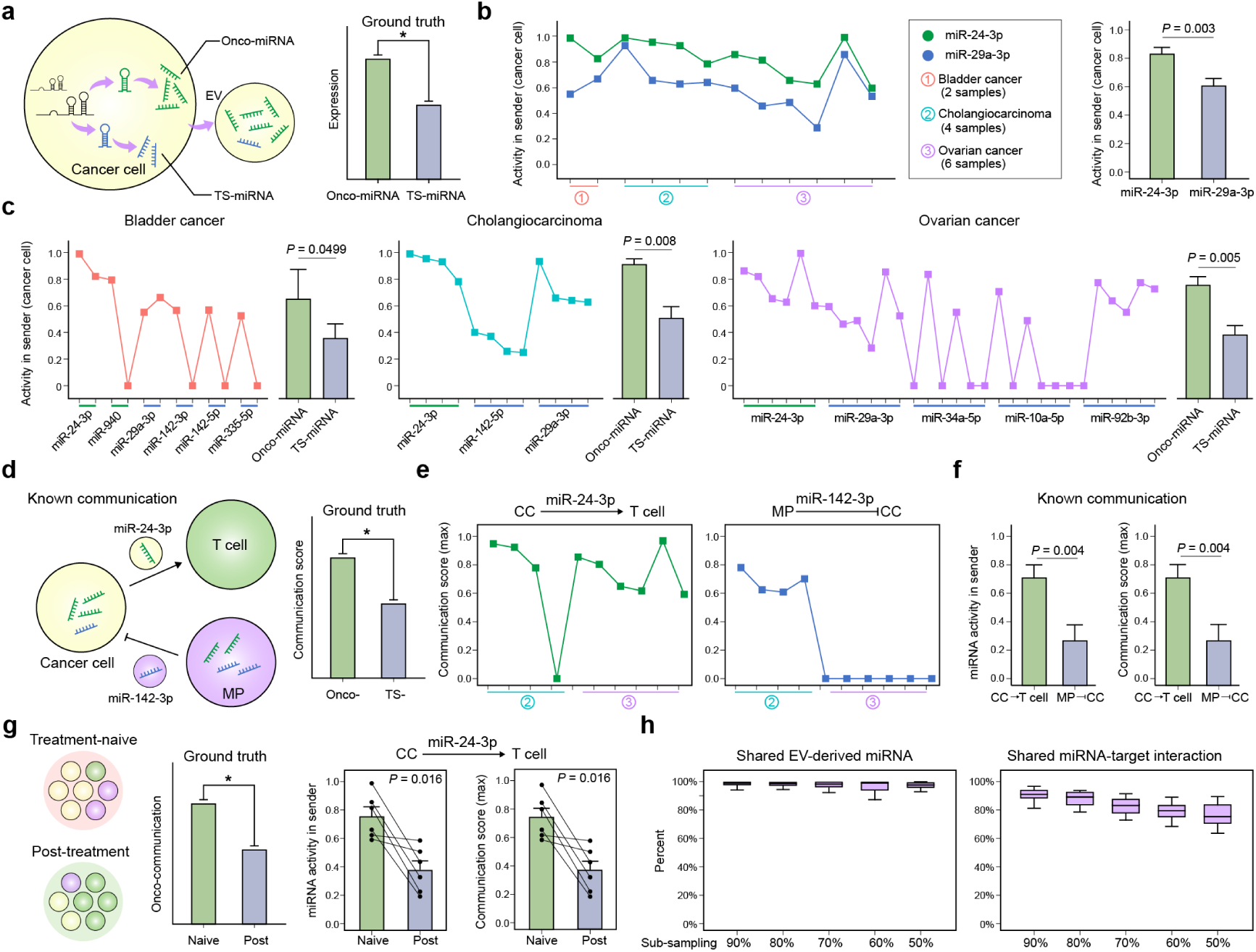
Prediction of miRTalk consistent with the literature evidence on real datasets. **(a)** Hypothesis regarding that the expression of oncogenic miRNAs is expected to surpass that of tumor suppressor (TS)-miRNAs within CCs without intervention. **(b)** Inferred activities and differential analyses of miR-24-3p and miR-29a-3p in CCs, represented across bladder cancer, cholangiocarcinoma, and ovarian cancer samples. **(c)** Detailed predictions of oncogenic miRNAs and TS-miRNAs in CCs among the bladder cancer, cholangiocarcinoma, and ovarian cancer samples. In the case of bladder cancer, oncogenic miRNAs include miR-24-3p and miR-940, while TS-miRNAs include miR-29a-3p, miR-142-3p, miR-142-5p, miR-335-5p. For cholangiocarcinoma, the oncogenic miRNA includes miR-24-3p, while TS-miRNAs include miR-142-5p and miR-29a-3p. In ovarian cancer, the oncogenic miRNA includes miR-24-3p, while TS-miRNAs include miR-29a-3p, miR-34-5p, miR-10a-5p, and miR-92b-3p. **(d)** Hypothesis postulating that the inferred oncogenic communication scores are expected to be higher than the TS-communication scores in CCs without intervention. **(e)** Inferred scores of cancer-T and MP-cancer communications mediated by the miR-24-3p and the miR-142-3p, respectively. CCs, cancer cells. **(f)** Difference of inferred miR-24-3p and the miR-142-3p activities as well as the maximum scores for the CC-T and MP-CC communications, respectively. **(g)** Hypothesis positing that the inferred oncogenic communication scores in tumors without intervention are expected to be higher than those observed with effective anti-cancer drug intervention. The Bar plot illustrates the differences of scores for the CC-T communication mediated by miR-24-3p at treatment-naive and post-treatment stages. **(h)** The percent of shared miRNAs and miRNA-target interactions with sub-sampling ratios of 90%, 80%, 70%, 60%, and 50%, in comparison to initial predictions. Significant differences between two groups were calculated with the one-sided Wilcoxon test.

T cells and macrophages (MPs) are involved with the initiation and advancement of neoplastic growths. For instance, exosomal miR-24-3p, a well-established oncogenic miRNA in cancer cells (CCs), is known to target and adversely affect the proliferative and differentiation capacities of T cells, consequently fostering tumor propagation and dissemination[36]. Concurrently, macrophages have been shown to secret exosomal miR-142-3p, which selectively targets and effectively suppresses the proliferation of CCs within the TME (**Fig. 3d**). Considering the disparate roles of miR-24-3p and miR-142-3p, we posited that, in the absence of intervention, the inferred scores for EV-derived miR-24-3p-mediated communication between CCs and T cells would surpass those of EV-derived miR-142-3p-mediated communication from macrophages to CCs within the TME[37]. With bladder cancer datasets lacking T cell data, we proceeded to assess miRTalk’s predictions using cholangiocarcinoma and ovarian cancer datasets. Interestingly, we noted elevated scores for EV-derived miR-24-3p-mediated CC-T communication in the majority of specimens, as inferred through miRTalk (**Fig. 3e**). In contrast, EV-derived miR-142-3p-mediated MP-CC communication was scarce within the ovarian TME, despite the observed presence of MIR142 that encodes miR-142-3p in the scRNA-seq data of ovarian cancer (**Supplementary Fig. S1i**). Consequently, both the oncogenic communication scores and oncogenic miRNA activities were significantly higher than their tumor suppressor counterparts (**Fig. 3f**), corroborating prior findings.

Furthermore, we reasoned that within the TME, scores for oncogenic miRNA-mediated CCC would experience notable reductions following effective therapeutic interventions. Hence, we employed miRTalk to infer oncogenic miRNA-mediated cancer cell-T cell communication by examining scRNA-seq datasets derived from treatment-naïve and post-treatment ovarian cancer patients (**Fig. 3g**). After including individuals with platinum-free intervals surpassing 50 days[21], we ascertained that post-treatment, all patients exhibited significantly diminished activities of oncogenic miR-24-3p originating from CCs. Correspondingly, the oncogenic miR-24-3p-mediated cancer cell-T cell communication scores substantially decreased, implying a restoration of T cell functionality attributable to reduced exosomal miR-24-3p secretions by CCs.

By conducting a stratified sampling procedure on the initial scRNA-seq data, we further investigated the efficacy of miRTalk inference across these authentic datasets. As a consequence, although a marginally lessened percentage of shared miRNA-target interactions was observed concomitant with the diminishing sub-sampling ratios, miRTalk persistently discerned a significant proportion of shared EV-derived miRNAs at sub-sampling ratios of 90%, 80%, 70%, 60%, and 50%, in juxtaposition with the initial predictions (**Fig. 3h**). These findings underscore the high accuracy and robustness of miRTalk in elucidating the biologically pertinent CCC mediated by EV-derived miRNAs.

### Metabolic modulation of glioblastoma cells on the TME sensed by nonmalignant cells

We first applied miRTalk to a human glioblastoma (GBM) scRNA-seq dataset to explore the CCC mediated by EV-derived miRNAs within the TME, comprising malignant glioblastoma cells, astrocytes (Astro), oligodendrocytes (Oligo), endothelial cells (Endo), oligodendrocyte precursor cells (OPC), neurons, and MPs[38] (**Fig. 4a**). As a consequence, intimate crosstalk within the GBM TME was detected, with malignant cells and MPs primarily functioning as prominent sources of miRNA secretion via EVs to other cells, rather than as receivers (**Fig. 4b, Supplementary Fig. S2a**). This finding is compatible with the understanding that malignant cells modulate the TME towards tumor promotion through the secretion of various EV types[39]. In harmony with this, numerous miRNAs such as miR-26a-5p, miR-221/222-3p, and miR-221/222-5p were identified in malignant cells and MPs (**Fig. 4c**). Prior research has reported that exosomal miR-26a derived from glioma stem cells can stimulate angiogenesis in Endo via the activation of the PI3K-AKT signaling pathway[40], corroborating the inferred communication from malignant cells to Endo through miR-26a-5p. This is further evidenced by the elevated expression of *MIR26A* and *MIR26A2* that encode miR-26a-5p and heightened gene set variation analysis (GSVA) scores for the PI3K/AKT signaling pathway in Endo (**Supplementary Fig. S2b-c**). Additionally, the highly expressed miR-221-3p presented a unique distribution within MPs (**Supplementary Fig. S2d**), acting as a critical mediator for communication between malignant cells and other cells (**Supplementary Fig. S2e**), aligning with the observations that miR-221-3p transfer via MP-derived EVs enhances GBM progression[41].

**Fig. 4.**
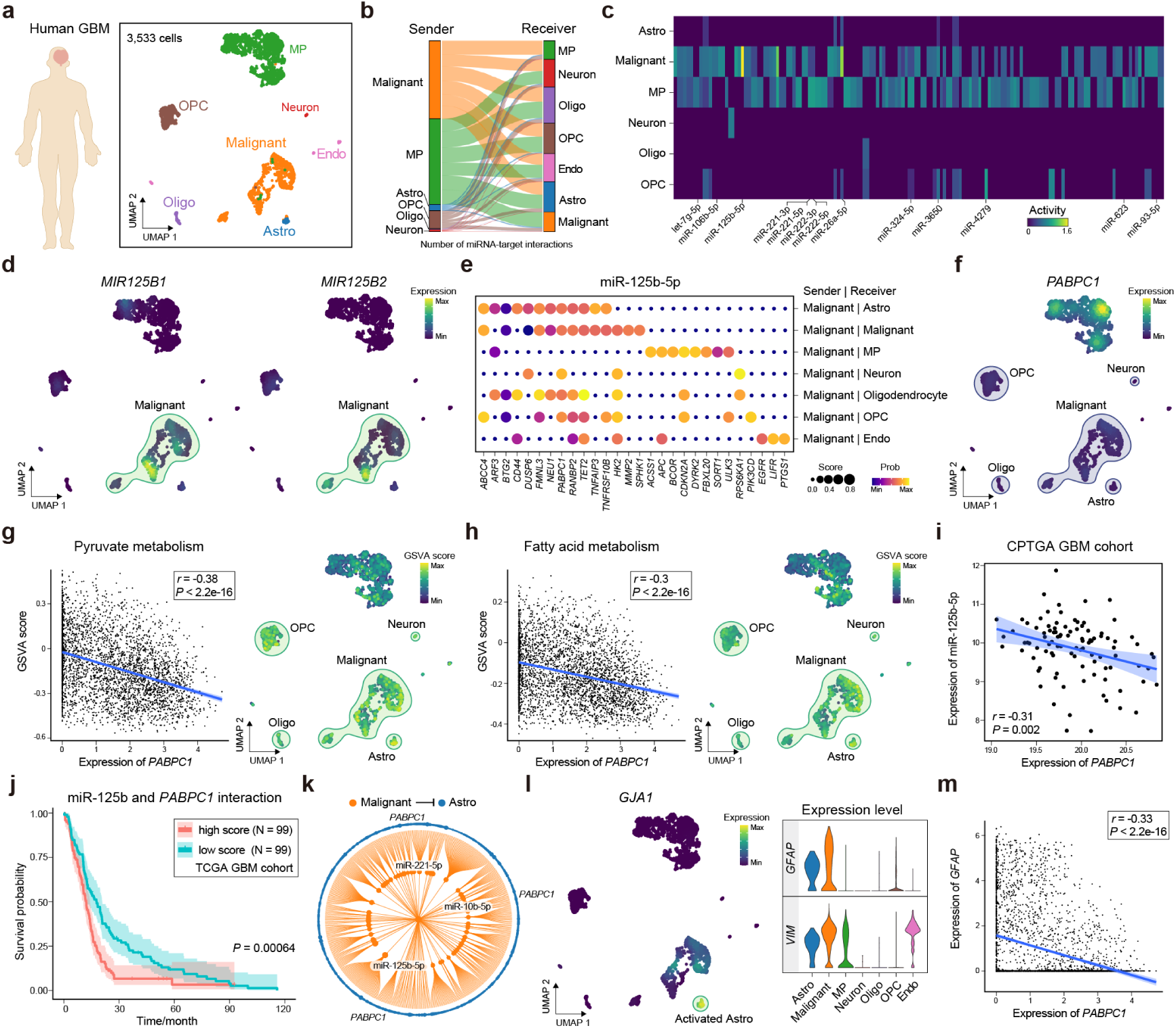
Metabolic modulation of glioblastoma cells on the TME sensed by nonmalignant cells. **(a)** The human glioblastoma (GBM) scRNA-seq dataset comprising 3,533 cells, encompassing malignant cells, astrocytes (Astro), oligodendrocytes (Oligo), endothelial cells (Endo), Neurons, macrophages (MP), oligodendrocyte precursor cells (OPC). **(b)** EV-derived miRNA-mediated cell-cell communications inferred by miRTalk, delineating the number of miRNA-target interactions amid pairwise cell types in GBM. **(c)** Heatmap portraying the activities of inferred EV-derived miRNAs among senders, i.e., Astro, malignant cells, MP, neurons, Oligo, and OPC. **(d)** Expression of *MIR125B1* and *MIR125B2* that encode the miR-125b-5p shown in the UMAP plot. **(e)** Autocrine and paracrine communication scores and probabilities of inferred miR-125b-5p-target interactions from malignant cells to other cell types. **(f)** UMAP plot displaying the expression of *PABPC1*. **(g-h)** Inverse correlation between the GSVA score of gene signatures and the expression of *PABPC1* analyzed with the Pearson’s coefficient. The gene signatures include pyruvate metabolism and fatty acid metabolism collected from the MSigDB. **(i)** Verification of the negative association between the expression of miR-125b-5p and PABPC1 within the CPTGA GBM cohort. **(j)** Survival analysis for patients respectively with high and low interaction scores between miR-125b and *PABPC1* in the TCGA GBM cohort. **(k)** Inferred miRNA-target interactions from malignant cells to Astro. **(l)** Expression of *GJA1*, *GFAP*, and *VIM*, signifying the activated Astro modulated by malignant cells. **(m)** Negative correlation between the expression of *PABPC1* and *GFAP*, assessed utilizing the Pearson’s coefficient.

Notably, our observations revealed that miR-125b-5p originating from malignant cells demonstrated the most pronounced activity among the various sender cell types, displaying a predominant prevalence of *MIR125B1* and *MIR125B2* which encode miR-125b-5p (**Fig. 4d**). Previous reports have indicated a significant correlation between elevated miR-125b levels and reduced survival rates in GBM patients[42], thus highlighting its critical role in modulating the malignancy and prognostication of GBM. Consequently, we investigated the mechanism of CCC between malignant cells and other cells within the TME, focusing particularly on miR-125b-5p. Through the secretion of miR-125b-5p via EVs, malignant GBM cells engaged in extensive interactions with neighboring cells through both autocrine and paracrine pathways (**Fig. 4e**). This results in the suppression of numerous target genes, such as *BTG2*, *TET2*, and *TNFRSF10B* among the corresponding receiver cells (**Supplementary Fig. 2f-g**). It is worth noting that *BTG2* has been extensively documented as a negative regulator of cell proliferation[43], while *TNFRSF10B* has been implicated in defense responses to tumor cells and apoptotic processes[44], thereby suggesting an enhancement of cell growth signaling within the GBM TME. As a methylcytosine dioxygenase responsible for maintaining normal DNA methylation processes, diminished *TET2* expression has been linked to unfavorable outcomes in multiple neoplasms[45], further corroborating previous findings.

Furthermore, we observed that *PABPC1* was targeted by EV-derived miR-125b-5p across various contiguous cells, including Astro, malignant cells, Oligo, OPC, and neurons (**Fig. 4f**). As a locus encoding a poly(A) binding protein, *PABPC1* demonstrated a significantly negative correlation with metabolism-related biological processes, such as glycolysis, gluconeogenesis, as well as pyruvate and fatty acid metabolism, evidenced by the elevated signature scores within these recipient cells (**Fig. 4g-h, Supplementary Fig. S2h**). The augmented metabolic processes intertwined with tumor progression have been extensively delineated within the TME, implying that malignant GBM cells exert metabolic modulation on the TME sensed by non-malignant cells through the EV-derived miR-125b-5p-*PABPC1* interaction. To verify the crucial role of this interaction within the GBM TME, we obtained an independent GBM cohort from the Clinical Proteomic Tumor Analysis Consortium[46] (CPTAC), which comprised sequencing data for miRNAs and mRNAs from 99 GBM patients. Subsequently, we observed a substantially negative association between the expression of miR-125b-5p and *PABPC1*, as well as a similar relationship between the expression of MIR125B1/MIR125B2 and *PABPC1* (**Fig. 4i, Supplementary Fig. S2i-j**). Considering the evidence substantiating the relationship between downregulated *PABPC1* and increased malignancy[47], as well as reduced survival rates in gliomas[48], we further investigated the association between miR-125b-*PABPC1* interaction and the prognosis of GBM. To this end, we divided patients from The Cancer Genome Atlas[49] (TCGA) GBM cohort into two distinct groups, respectively characterized by high and low scores for miR-125b-*PABPC1* interaction. Remarkably, the group of GBM patients exhibiting elevated miR-125b expression and diminished *PABPC1* expression displayed a significantly abbreviated survival duration (**Fig. 4j**). In fact, the pervasive consensus in the scientific community acknowledges that the oncogenic transformation of the metabolic TME by tumors can establish an immunosuppressive milieu[50], thereby undermining the efficacy of the host’s anti-tumor immune response. This phenomenon could potentially explain the adverse prognosis associated with an intensified miR-125b-*PABPC1* interaction in GBM patients.

It is known that Astro constitute approximately half of the brain’s total cell population, and their pivotal role in GBM progression has been increasingly studied[39]. Upon activation in response to tumor-derived EVs, Astro can establish an immunologically advantageous environment for tumor cells, promoting further tumor growth and invasion[39]. In harmony with these findings, Astro displayed the most responsiveness towards miR-125b-5p emanating from malignant GBM cells, as evidenced by the highest aggregate communication scores between the two entities (**Supplementary Fig. S2k**). Alongside miR-125-5p, the well-established oncogenic miRNAs in GBM, miR-221-5, and miR-10b-5p, played a significant role in regulating Astro (**Fig. 4k**), which exhibited an activated phenotype marked by heightened expression of *GFAP*, *GJA1*, and *VIM* (**Fig. 4l, Supplementary Fig. S2l**). Intriguingly, a markedly inverse correlation was observed between *PABPC1* expression and signatures of activated Astro (**Fig. 4m, Supplementary Fig. S2m**), thereby suggesting potential roles for malignant GBM cell-derived miRNAs in orchestrating the activation of surrounding Astro within the TME.

### Characterization of signal transmissions forming the fibrogenic niche in kidney

A considerable corpus of empirical findings emphasizes the involvement of miRNAs in renal impairment and pathology. Consequently, a murine scRNA-seq dataset of kidney fibrosis induced by unilateral ureteral obstruction (UUO) was used for extrapolating EV-derived CCC amongst vascular smooth muscle cells (VSMCs), injured VSMCs, mesangial cells, pericytes, parietal epithelial cells (PECs), and myofibroblasts[51] (**Fig. 5a, Supplementary Fig. S3a**). As a result, intricate CCC networks governed by EV-derived miRNAs were inferred (**Supplementary Fig. S3b**), wherein numerous well-established miRNAs associated with renal fibrosis were identified[52, 53], including miR-23a-3p, miR-27a-3p, and miR-145-5p (**Fig. 5b**). Remarkably, injured VSMCs emerged as the most dynamic cellular cohort[54], secreting an array of EV-derived miRNAs to other populations. This underscored the pivotal role of injured VSMCs in modulating the fibrogenic microenvironment responses to UUO-induced damage.

**Fig. 5.**
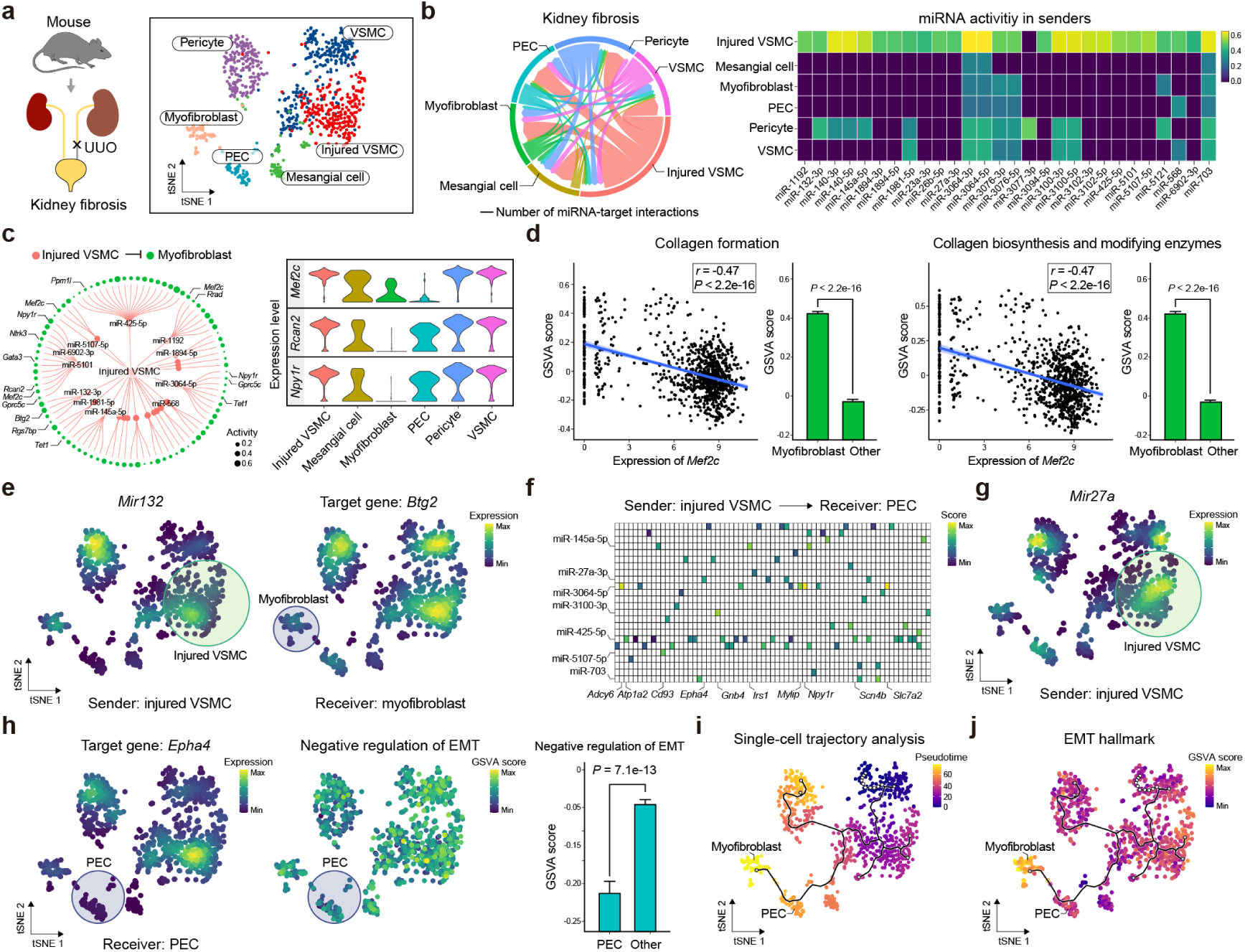
Characterization of signal transmissions forming the fibrogenic niche in kidney. **(a)** The mouse scRNA-seq dataset of the fibrotic kidney comprising vascular smooth muscle cells (VSMCs), injured VSMCs, mesangial cells, pericytes, parietal epithelial cells (PECs), and myofibroblasts. UUO, unilateral ureteral obstruction. **(b)** EV-derived miRNA-mediated cell-cell communications inferred by miRTalk, displaying the number of miRNA-target interactions between pairwise cell types in fibrotic kidney. Heatmap showing the activities of inferred EV-derived miRNAs for senders. **(c)** Inferred miRNA-target interactions from injured VSMCs to myofibroblasts with the violin plots depicting the expression of target genes, i.e., *Mef2c*, *Rcan2*, and *Npy1r*. **(d)** Negative correlation between the GSVA score of gene signatures and the expression of *Mef2c* analyzed with the Pearson’s coefficient. The gene signatures include collagen formation and collagen biosynthesis and modifying enzymes collected from the MSigDB. **(e)** UMAP plot showing the expression of *Mir132* and its target gene *Btg2*. **(f)** Injured VSMC-PEC communication scores of inferred miRNA-target interactions. **(g)** Expression of *Mir27a* in the UMAP plot. **(h)** UMAP plot about the expression of the target gene *Epha4* for miR-27a-3p and the GSVA score for the negative regulation of EMT. **(i)** Single-cell trajectory analysis with the monocle3 for the construction of the pseudotime trajectory. **(j)** GSVA score for the EMT hallmark collected from the MSigDB. Significant differences between two groups were calculated with the one-sided Wilcoxon test.

Renal fibrosis is typified by the undue accumulation and deposition of extracellular matrix (ECM). While a multitude of cell types can synthesize ECM, it is widely held that myofibroblasts constitute the primary cellular component contributing to fibrosis in afflicted kidneys[55]. In light of this, we explored the EV-derived miRNA-mediated interplay between injured VSMCs and myofibroblasts. Accompanied by EV-derived miRNAs, injured VSMCs exerted appreciable inhibitory regulation on a series of target genes, such as *Mef2c*, *Rcan2*, *Npy1r*, *Rgs7bp*, and *Gprc5c*, within myofibroblasts (**Fig. 5c, Supplementary Fig. S3c**). The majority of these target genes displayed a considerable negative association with the fibrogenic collagen-formation pathway contributing to ECM production (**Supplementary Fig. S3d**). For instance, *Mef2c* in myofibroblasts—concomitantly targeted by injured VSMC-derived miR-1192, miR-5101, and miR-5107-5p—demonstrated substantial negative correlations between its expression and the GSVA scores for both the collagen formation pathway and the pathway related to collagen biosynthesis and modifying enzymes. Moreover, significantly elevated GSVA scores were observed in myofibroblasts compared to alternative cellular populations (**Fig. 5d, Supplementary Fig. S3e**), potentially insinuating the essential function of *Mef2c* in responding to the inhibitory cues emanating from injured VSMCs. Additionally, the highly expressed miRNA-132 bolsters the development of renal fibrosis by selectively enhancing myofibroblast proliferation—another major event in establishing the fibrogenic niche within the kidney[56, 57]. Aligning with this, the injured VSMC-myofibroblast crosstalk facilitated by miR-132-3p was significantly discerned, coupled with the corresponding downregulated target gene *Btg2*, a known contributor to the negative regulation of cell population proliferation[43].

As an essential component of resident glomerular cells, PECs have garnered considerable attention in recent years, fostering more comprehensive understanding and delineation of their potential roles in renal pathology[58]. In this context, we studied the EV-derived miRNA-mediated crosstalk between injured VSMCs and PECs, yielding a multifaceted miRNA-target interaction network (**Fig. 5f**). In concordance with a recent investigation stating that miR-27a-3p silencing ameliorated effect on renal fibrosis[59], we observed the elevated expression of miR-27a-3p within the fibrogenic niche, particularly in injured VSMCs (**Fig. 5g**), concurrent with the preeminent interaction of miR-27a-3p and *Epha4*, specifically identified between injured VSMCs and PECs (**Supplementary Fig. S3f-g**). Considering *Epha4*’s role in the negative regulation of epithelial-to-mesenchymal transition[60] (EMT), the corresponding GSVA scores in PECs were found to be significantly lower than those of other cells, thereby implying an augmented EMT activity in PECs (**Fig. 5h**). As a result, a single-cell trajectory analysis employing monocle3 was used to dissect cellular decisions for PECs, culminating in the reconstruction of a branching trajectory from the root to the end states (**Fig. 5i**). In alignment with this, the EMT scores intensified along the pseudotime axis, transitioning from PECs to myofibroblasts, which indicated that PECs contribute to the origin of myofibroblasts upon communication with injured VSMCs. These findings potentially elucidated the communicative mechanisms that underpin signal transmissions between injured VSMCs and myofibroblasts, thereby promoting the establishment of the fibrogenic niche in kidney.

### Identification of the granulocyte-hepatocyte communication discriminating between the normal and fatty liver transplantation

To ascertain the efficacy of miRTalk in scRNA-seq datasets under disparate conditions, we employed it on our previously published data encompassing normal and high-fat diet transplanted rat livers[61]. Our objective was to contrast the EV-derived miRNA-mediated CCC within normal and fatty liver transplantation (LT) settings. Our analysis categorized a total of 23,675 cells as granulocytes, MPs, monocytes, DCs, hepatocytes, NK cells, Kupffer cells, T cells, Endo, and B cells, derived from six transplanted liver samples procured from three normal and three high-fat diet rats (**Fig. 6a, Supplementary Fig. S4a**). In light of miRNA inference, an abundance of EV-derived miRNA-mediated CCC transpired, particularly between hepatic parenchymal and non-parenchymal cells, with a more intricate communication network discerned for fatty livers (**Fig. 6b**). This was substantiated by the elevated score of miRNA-target associations observed in fatty transplanted livers relative to normal transplanted livers (**Supplementary Fig. S4b**). As the predominant cell population in the liver, hepatocytes exhibited an inclination to receive EV-derived miRNAs from hepatic non-parenchymal cells, with a proclivity against secretion after LT (**Fig. 6c**). Notably, hepatocytes within fatty transplanted livers had a greater activity of EV-derived miRNAs and score of miRNA-target interactions in contrast to normal transplanted livers (**Fig. 6d, Supplementary Fig. S4c**), insinuating that enhanced communication from hepatic non-parenchymal cells to hepatocytes was orchestrated by the steatotic microenvironment.

**Fig. 6.**
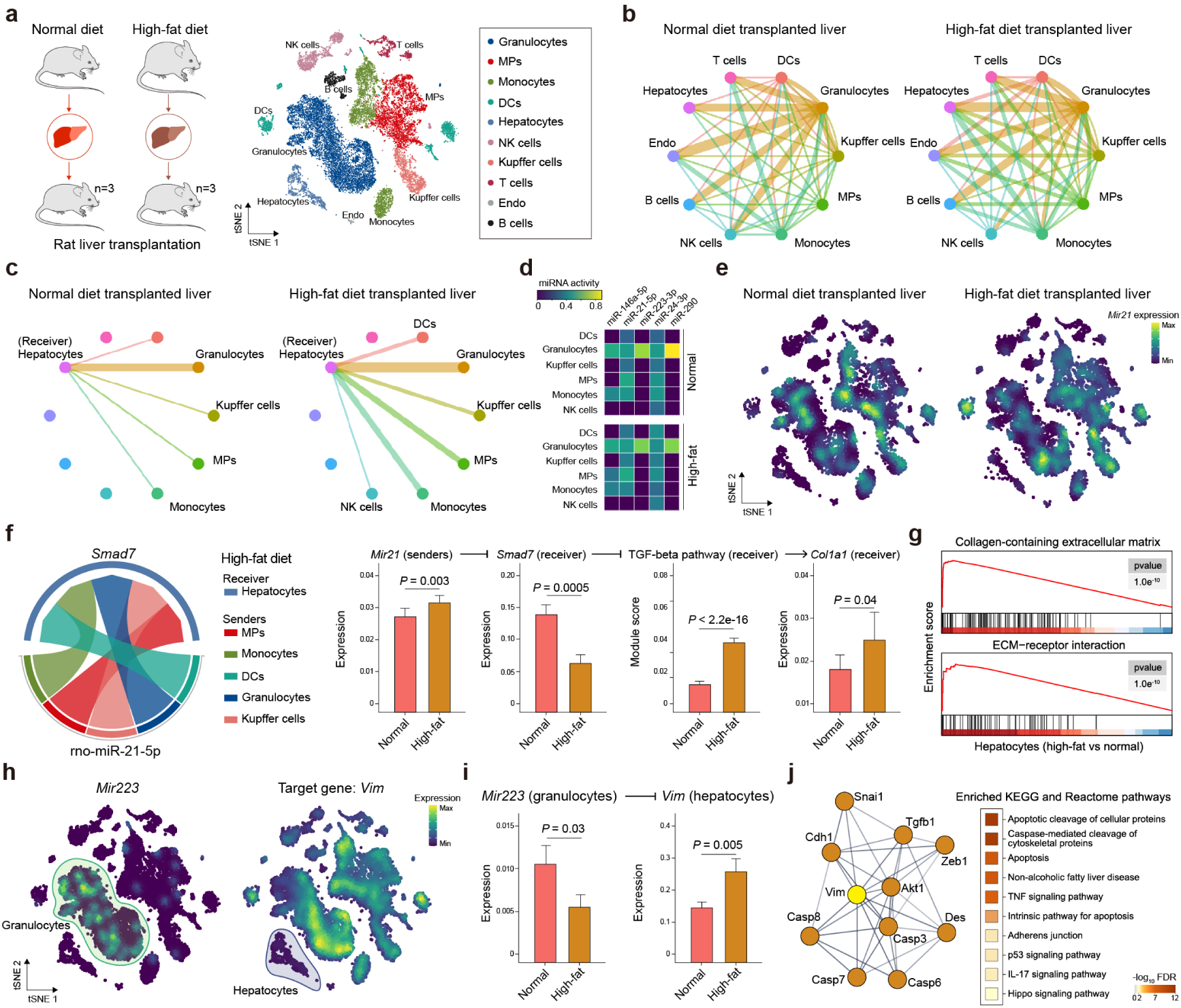
Identification of the granulocyte-hepatocyte communication discriminating between the normal and fatty liver transplantation. **(a)** The rat scRNA-seq dataset of the normal and high-fat diet transplanted livers including granulocytes, MPs, monocytes, DCs, hepatocytes, NK cells, Kupffer cells, T cells, Endo, and B cells. **(b)** EV-derived miRNA-mediated cell-cell communications inferred by miRTalk, displaying the number of miRNA-target interactions between pairwise cell types in normal and high-fat diet livers. **(c)** Number of miRNA-target interactions with EV-derived miRNAs sent from immune cells to hepatocytes. **(d)** Heatmap showing the activities of inferred EV-derived miRNAs for senders. **(e)** UMAP plot of the gene expression for *Mir21* in normal and high-fat diet livers. **(f)** The interaction between miR-21-5p and its target gene *Smad7* that mediates the communication from other cells to hepatocytes in fatty livers. Significant differences between two groups were calculated with the one-sided Wilcoxon test and Welch test. **(g)** Gene set enrichment analysis (GSEA) with gene sets from MSigDB by comparing the differentially expressed genes (DEGs) of hepatocytes collected from normal and fatty transplanted livers. **(h)** Expression of *Mir223* in granulocytes and Vim in hepatocytes. **(i)** Comparison of the expression *Mir223* in granulocytes and Vim in hepatocytes between the normal and high-fat diet livers with one-sided Wilcoxon test. **(j)** Protein-protein interaction network analysis of Vim and the enriched KEGG pathways and Reactome pathways using the STRING web tool.

Specifically, we identified a pervasive presence of miR-21-5p within immune cells inhabiting both normal and fatty environments, exerting a marked inhibitory effect on *Smad7* expression (**Fig. 6e, Supplementary Fig. S4d**) – a molecule known to negatively modulate the TGF-beta signaling cascade[62]. Intriguingly, while the association between miR-21-5p and its target gene, *Smad7*, remained unaltered across the distinct hepatic milieus, we observed a stark contrast in the relative abundance of these molecules (**Fig. 6f, Supplementary Fig. S4e**). In the context of high-fat diet livers, sender cells (MPs, monocytes, DCs, granulocytes, and Kupffer cells) exhibited a pronounced amplification in miR-21-5p expression as opposed to their counterparts in normal diet livers. Concurrently, *Smad7* expression within recipient hepatocytes was substantially diminished in high-fat livers – an observation that corroborates the elevated miR-21-5p expression in this context[63]. As anticipated, hepatocytes residing in high-fat diet environments displayed a significantly accentuated TGF-beta pathway module score relative to those within normal diet livers. Moreover, elevated expression of *Col1a1*—a gene encoding the pro-alpha1 chains of type I collagen implicated in hepatic fibrosis onset—was detected in high-fat diet hepatocytes (**Fig. 6f**). Comprehensive gene set enrichment analysis (GSEA)[64] employing differentially expressed genes (DEGs) and the molecular signature database[65] (MSigDB) elucidates the heightened fibrotic response in high-fat livers via the significant enrichment of collagen-containing matrix and ECM-receptor interaction gene sets (**Fig. 6g**). Collectively, these findings revealed that steatotic perturbations in CCC mediated by EV-derived miRNAs play a vital role in shaping the hepatic susceptibility to fibrosis under high-fat diet conditions.

Intriguingly, granulocytes emerged as the predominant regulators of hepatocyte target genes in both normal and steatotic transplanted livers, through the action of EV-derived miRNAs (**Fig. 6d**). Given their status as the most abundant cellular population within the hepatic milieu, granulocytes have been implicated in critical processes such as ischemia-reperfusion injury and liver fibrosis[66]. Notably, miR-223-3p expressed in granulocytes exerted a substantial inhibitory influence on hepatocytes by downregulating its target gene named *Vim* – a mediator implicated in ischemia-reperfusion injury (**Fig. 6h**). In comparison to normal diet transplanted livers, high-fat diet ones exhibited a significant downregulation of *Mir223* in granulocytes and upregulation of *Vim* in hepatocytes, consistent with the evidence demonstrating the amelioration of neutrophil-derived miR-223 on nonalcoholic steatohepatitis by communicating with hepatocytes[6] (**Fig. 6i**). Focusing on the protein-protein interaction network centered on Vim, we identified a significant enrichment of apoptosis-related and TNF-signaling pathways (**Fig. 6j**). This suggests that hepatocytes within high-fat diet livers experience heightened apoptotic signals as a consequence of interactions between granulocyte-derived miR-223-3p and its target gene *Vim*. In contrast, hepatocytes within normal diet livers following LT continued to maintain normal metabolic functions, as evidenced by the significant enrichment of metabolism-related pathways such as steroid metabolic process, fatty acid metabolic process, and lipid metabolism (**Supplementary Fig. S4f**). These findings underscore the disruption of hepatic metabolic function through EV-derived miRNA-mediated communication between hepatocytes and surrounding cells, such as granulocytes, within the steatotic microenvironment.

## Discussion

In this study, we introduced a pioneering computational approach called miRTalk to infer EV-derived miRNA-mediated CCC employing scRNA-seq data. Incorporating a probabilistic model and a curated database—miRTalkDB—containing EV-derived miRNA-target interactions for Homo sapiens, Mus musculus, and Rattus norvegicus, this method attests its novelty. Assessment against both simulated and real-world datasets substantiated the exceptional accuracy and robustness of miRTalk. Moreover, analyzing three independent scRNA-seq datasets using miRTalk unveiled profound communicative mechanisms within the diseased contexts.

CCC, also known as cell-cell interaction, is an essential feature in multicellular organisms [67]. With the widespread adoption of scRNA-seq data, it is customary to combine the abundance of ligands and receptors to infer signal transmissions from sender to receiver cells by reasoning that highly co-expressed ligands and receptors are inclined to facilitate ligand-receptor interaction (LRI)-dependent CCC. Examples include CellChat[17], CellPhoneDB[16], NICHES[18], ICELLNET[19], and scTenifoldXct[68]. Another approach utilizes downstream targets stimulated by LRIs in receiver cells to enhance the accuracy of inferred CCC, primarily based on highly co-expressed ligands, receptors, and targets, such as NicheNet[69], CytoTalk[70], CellCall[71], and scMLnet[72]. Furthermore, advancements in spatially resolved transcriptomic methodologies enable the inference of spatially resolved juxtacrine and paracrine communication, reasoning that cells are more likely to engage in communication if they are spatially proximal. Consequently, several computational methods has emerged to infer cell-cell communication underlying normal and diseased tissues with spatial structures, such as SpaTalk[73], Giotto[74], SpaOTsc[75], spaCI[76], cell2cell[77], COMMOT[78], CCPLS[79], and SpaCET[80].

Recently, Zheng et al. pioneered the development of a computational algorithm known as MEBOCOST to infer CCC mediated by metabolites, including lipids, and their corresponding sensor proteins for scRNA-seq data[81]. Likewise, MEBOCOST posited that metabolite-driven CCC arises when metabolite enzymes and sensors are highly expressed in sender and receiver cells. Given that neurons can extend axons and dendrites across considerable distances to form synapses and communicate primarily through neurotransmitters, Zhao et al. consequently established NeuronChat, a tailored method for inferring neural-specific CCC utilizing scRNA-seq data based on the coordinated expressions of curated ligands and their targets[82].

Indeed, beyond the CCC facilitated by LRIs, spatial contact, metabolites, and neurotransmitters aforementioned, EV-derived miRNAs also embody critical chemical signals mediating short-range and long-range CCC. Contrasting the hypothesis present in existing CCC inference approaches, miRTalk postulates that CCC transpires when miRNA expression levels are elevated in sender cells, concurrently with diminished expression of the associated target genes in receiver cells. Despite the distinct principles employed in miRTalk and existing methods, the excellent performance achieved by miRTalk as well as other methods in predicting positive and negative samples generated via the same pipeline substantiated the reliability of miRTalk in inferring EV-derived miRNA-mediated CCC.

To ensure the accurate inference of CCC within a specific context, it is a common practice to construct a tailored foundational database in the development of a novel inference method. For instance, CellPhoneDB integrates the subunit architecture of both ligands and receptors, representing heteromeric complexes, while CellChatDB encompasses various cofactor types, such as soluble agonists and antagonists, as well as co-stimulatory and co-inhibitory membrane-bound receptors. To accommodate the inference of metabolite-dependent intercellular communication and neurotransmitter-dependent neuron-neuron interactions, MEBOCOST and NeuronChat curated databases of metabolite-sensor partnerships and ligand-target interactions, respectively. For the accurate inference of spatially resolved intercellular communication, the SpaTalk method relies on the foundational CellTalkDB[83] and transcription factor (TF)-target interaction database. Consequently, we curated an EV-derived miRNA-target database, termed miRTalkDB, by incorporating experimentally verified information on miRNA-target interactions.

For miRTalk, the inferred EV-derived miRNAs predominantly rely upon the extent of miRNA coverage in scRNA-seq data. Despite the coverage of miRNAs in contemporary datasets produced by prevalent scRNA-seq methodologies being rather sparse, miRTalk successfully discerned an abundance of EV-derived miRNAs from the scRNA-seq data. Fortuitously, the recently reported snRandom-seq demonstrated enhanced coverage of non-coding genetic elements, such as miRNAs, within their scRNA-seq data by procuring full-length total RNAs via random primers[84]. Moreover, Ji et al. proposed an efficient and straightforward computational framework to profile miRNAs from scRNA-seq data named PPMS[85], which potentially offers supplementary abundance of miRNAs of interest. The ongoing expansion of scRNA-seq data, characterized by augmented accuracy, abundance, and coverage of miRNAs through such innovative technologies and computational instruments, will undoubtedly facilitate the practicality of miRTalk across a wide spectrum of applications in the fields of biology and biomedicine.

## Conclusions

In summary, miRTalk represents the pioneering method for inferring EV-derived miRNA-mediated CCC from scRNA-seq data by incorporating a curated database encompassing EV-derived miRNA-target associations for Homo sapiens, Mus musculus, and Rattus norvegicus, offering new insights into the intercellular dynamics underlying biological processes.

## Methods

### Datasets

The scRNA-seq dataset of cholangiocarcinoma in humans was directly accessed from Gene Expression Omnibus (GEO) repository: GSE138709[10]. The human scRNA-seq datasets of bladder cancer, glioblastoma multiforme, and ovarian cancer were curated from the TISCH2 database (http://tisch.comp-genomics.org/) with the accession numbers GSE130001[20], GSE84465[38], and GSE165897[21], respectively. The murine scRNA-seq data of normal and fibrotic kidneys was procured from the Zenodo data archive (https://zenodo.org/record/4059315). The previously published scRNA-seq data of normal and fatty liver transplantation was deposited in the Genome Sequence Archive (GSA) in the National Genomics Data Center (NGDC) under the accession number CRA004061[86]. The glioblastoma multiforme datasets from The Cancer Genome Atlas[49] (TCGA) and the Clinical Proteomic Tumor Analysis Consortium[46] (CPTAC) incorporating the miRNA and mRNA expression profiles were accessed from the cBioPortal (https://www.cbioportal.org/datasets) web resource[87].

### Data processing

For the analyses of human ovarian cancer investigations, patient participants with platinum-free interval exceeding 50 days and corresponding peritoneum and omentum samples containing at a minimum 100 epithelial ovarian cancer cells in the scRNA-seq data were retained. The nomenclature of genes was rectified in accordance with the National Center for Biotechnology Information gene database (https://www.ncbi.nlm.nih.gov/gene/), in which unmatched and duplicated genes were removed. Normalized data were used by the log-normalization procedure of Seurat[88] for identifying the highly variable genes in the miRTalk analysis pipeline.

### Construction of miRTalkDB

The compendiums of Homo sapiens, murine, and Rattus norvegicus miRNAs were amassed from the well-known database named miRTarBase[14] (https://mirtarbase.cuhk.edu.cn/), which is an experimentally verified catalog of microRNA-target interaction networks. EV-derived miRNAs were filtered by examining their presence in the following established repositories: (1) ExoCarta[11] (http://exocarta.org/), an exosome database furnishing the content discerned in exosomes across various species; (2) Vesiclepedia[12] (http://microvesicles.org/), a meticulously curated repository of molecular information identified in distinct classes of EVs, such as ectosomes, microvesicles, exosomes, or apoptotic bodies; (3) EVmiRNA[13] (http://bioinfo.life.hust.edu.cn/EVmiRNA#!/), a database of miRNA expression patterns in EVs. For miRNAs not present within these databases, we conducted text mining within PubMed employing the use of critical keywords: “[miRNA name or aliases] AND (extracellular vesicles)”, followed by manual validation to discern corroborating evidence demonstrating the presence of relevant miRNAs in EVs. By integrating the EV-derived miRNA dataset and the empirically substantiated miRNA-target interactions, we curated the miRTalkDB as the foundational database to infer EV-derived miRNA-mediated CCC.

### Algorithm of miRTalk

The miRTalk model is composed of two principal components: identification of miRNA and target genes as well as enrichment of miRNA-target interactions. The former focuses on filtering the widely expressed miRNAs and the potential target genes with significant variability in expression regarding EV-derived miRNAs, while the latter aims to infer the EV-derived miRNA-target interactions that facilitate CCC from senders to receivers.

#### Identification of miRNA and target genes

To ensure that the inferred target genes in receiver cells accurately reflect the inhibitory effect of the EV-derived miRNA originating from sender cells, we initially endeavored to identify the highly variable target genes within a given scRNA-seq data matrix *G*[*g* × *c*] (*g* genes and *c* cells) with *k* cell types. Let *G*_*t*_ represent the highly variable target gene set, which can be calculated by the following function:

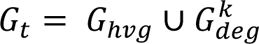

Herein, *G*_*hvg*_ refers to the highly variable genes (HVGs) procured from *G* utilizing the local polynomial regression (loess) and the variance stabilizing transformation method in Seurat. Concurrently, 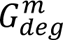 denotes the combined differentially expressed genes (DEGs) for *k* cell types, employing the Wilcoxon Rank-Sum (WRS) test, along with a minimal log-scale fold change of 0.25 in conjunction with a minimum of 10% expression in cells. Additionally, the widely expressed miRNAs were filtered with a minimum of 10 expressed cells for a given sender cell type.

#### Enrichment of miRNA-target interactions

To enhance the enrichment of miRNA-target interactions in the context of sender-receiver communication, we propose a filtering rule that considers the expression levels of miRNA and target genes in sender and receiver cells. Specifically, for a given sender data matrix *S*[*g* × *s*] (*g* genes, *n* miRNA genes, and *s* cells) and a receiver data matrix *R*[*g* × *r*] (*g* genes, *n* miRNA genes, and *r* cells), we assign the *i*^*th*^ (*i* ≤ *n*) miRNA gene expression value in a given sender cell to φ, and the *j*^*th*^ (*j* ≤ *g*) corresponding target gene expression value in a given receiver cell to τ, according to the miRNA-target pairs recorded in miRTalkDB. To enrich the miRNA-target interactions that potentially mediate the sender-receiver communication, we filter out the communication events where the miRNA gene in sender cells is highly expressed and the corresponding target gene in receiver cells is lowly expressed. To this end, we proposed a probabilistic model that deduces the miRNA-target interacting probability *c* with the following function:

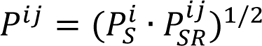

where 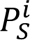 represents the probability of highly expressed miRNA in sender cells and defined as:

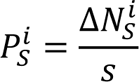

Here, 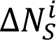 is the number of sender cells with higher miRNA expression than the average expression 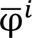 of the same miRNA in other cells. 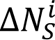 can be calculated using the following formula:

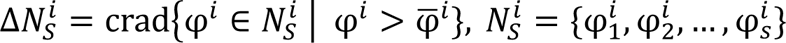

In terms of 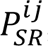, it represents the probability that the lowly expressed target gene in receiver cells is targeted by the EV-derived miRNA from sender cells, which can be determined as follows:

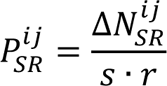

where 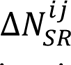 denotes the number of sender-receiver pairs with the miRNA gene expression in receiver cells 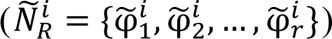 lower than that in sender cells 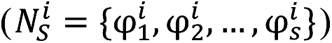 and the expression of target genes in receiver cells 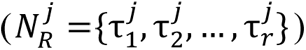 lower than the average expression 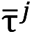 in other cells. This can be expressed as:

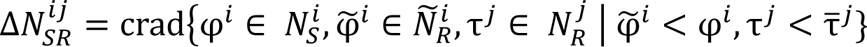

### Inference of miRNA and target activity

Considering that some mature miRNAs are encoded by multiple miRNA genes, and the sparse abundance of miRNA gene expression in scRNA-seq data matrix, we employed an aggregated function to infer the miRNA activity within each cell type. Let *m* be the number of the corresponding miRNA genes for a given miRNA; its activity, **A*_miR_*, can be written as:

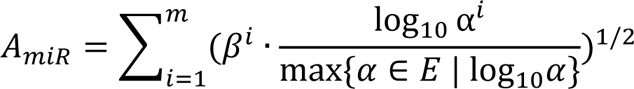

where *E* = {α^1^, α^2^, …, α^*n*^} represents the expressed percent of *n* miRNA genes among sender cells. For a given miRNA gene, α^*i*^ and β^*i*^ can be determined by the following functions:

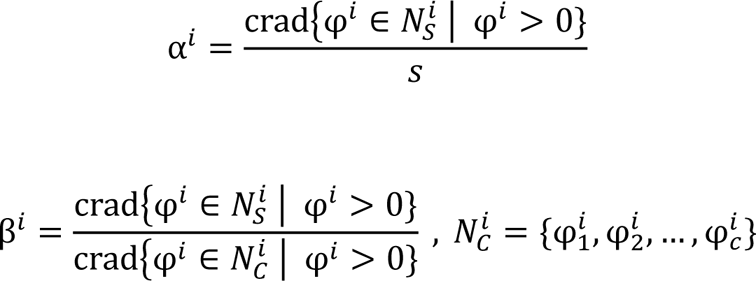

To ascertain the target gene activity in receiver cells, let *T* = {γ^1^, γ^2^, …, γ*^g^*} be the average expression of *g* genes among receiver cells, which can be determined via the following equation:

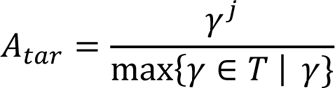

Upon determining miRNA and target activity, we integrated a miRNA-target interaction score **S*_*mii*R−tar_* and probability *P_*mii*R−tar_* for a given mature miRNA in sender cells and its corresponding target gene in receiver cells, expressed as:

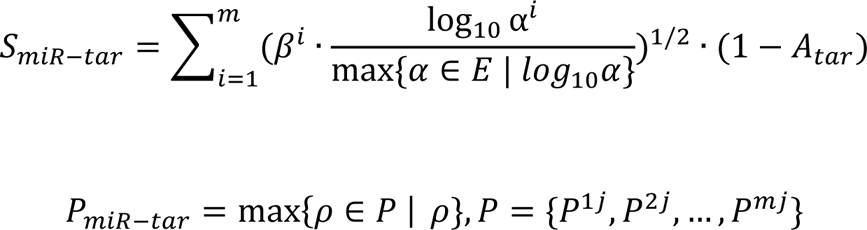

### Simulation of scRNA-seq data for benchmarking

The ESCO[15] R package was utilized to simulate the benchmarked datasets through variegating the gene and cell number. By setting parameters such as ‘nGenes = 1,000’, ‘nCells = 1,000’, ‘group.prob = c(0.5, 0.5) ‘, ‘deall.prob = 0.3’, ‘de.prob = c(0.5, 0.5)’, ‘de.facLoc = c(1.9, 2.5)’, ‘withcorr = TRUE’, and ‘trials = 1’, the initially generated scRNA-seq data matrix of 1,000 gene and cell pairings were derived, which is composed of a sender cell type containing 510 cells, and a receiver cell type containing 490 cells. For testing miRTalk, 1,000 genes were bifurcated into 500 miRNA genes matching with the remaining 500 target genes. Negative correlations were filtered out to form positive samples, where the miRNA genes in sender cells were significantly highly expressed, paired with the target genes in receiver cells that were significantly lowly expressed. In contrast, negative samples arose from interactions with the significantly lowly expressed miRNA genes in sender cells, paired with significantly highly expressed target genes in receiver cells. For testing other LR-based inference methods, the 1,000 genes were assigned into 500 ligand genes matched with the remaining 500 receptor genes. Positive samples were determined on the significantly highly expressed ligand genes in sender cells, paired with receptor genes in receiver cells that were both significantly highly expressed. Negative samples, on the other hand, arose from interactions with both significantly lowly expressed ligand genes in sender cells and receptor genes in receiver cells.

### Comparison with other methods

In order to conduct a thorough comparison of various CCC inference methods, our study utilized the simulated LR database that contained positively correlated LR interactions as supplement to the default database in each method, as described above. Through the variation of key parameters such as gene number (1,000, 800, 600, 400, and 200), ligands/receptors coverage (50%, 40%, 30%, 20%, and 10%), cell number (1,000, 800, 600, 400, and 200), and sender/receiver ratio (8:2, 6:4, 5:5, 4:6, and 2:8), we simulated 20 different datasets under four distinct contexts. This allowed us to test the performance of various methods while ensuring that we had solid ground truth for evaluation purposes. In practice, simulated data was transformed into the h5ad format with the SeuratDisk R package to run CellPhoneDB with the statistical analysis by default parameters, where the probability was determined by subtracting p-value from 1. For running pipelines of CellChat, NICHES, and ICELLNET, we used the simulated data directly to infer the LR interactions using default parameters. When using NICHES, we calculated the average score for a given LR interaction among all sender-receiver pairs as the probability. Additionally, for ICELLNET, we used the score divided by 100 to evaluate the probability of LR interactions. AUROC and AUPRC were calculated to evaluate the performance of predictions with the precrec[89] R package.

### Bulk data analysis

The Pearson correlation coefficient was calculated between the miR-125b-5p and its target gene *PABPC1* with the correlation R package. In order to investigate the relationship between the miR-125b-5p-*PABPC1* interaction and overall survival (OS) of patients with GBM, we applied the miRTalk algorithm to calculate the interaction score (*IS*) using the following formula:

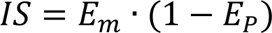

where *E*_*m*_ and *E*_*p*_ respectively represent the expression levels of miR-125b and *PABPC1*. After calculating the *IS* for each patient, we grouped the GBM patients into two categories based on their high or low score and analyzed their survival using the survival and survminer R packages with default parameters.

### Pathway and biological process enrichment

The Metascape web tool (https://metascape.org/) was used to carry out the enrichment of pathways and biological processes[90], wherein the top 100 highly expressed genes were chosen according to the fold change of the average gene expression. The GSVA[91] R package and the “AddModuleScore” function in Seurat were utilized to compute the module score of pathways, GO biological processes, and known hallmarks. To obtain the signatures, the Molecular Signatures Database[65] (MSigDB, http://www.gsea-msigdb.org/gsea/msigdb) was accessed through msigdbr. GSEA was conducted by clusterprofiler[64] with the ordered gene list and MSigDB. The STRING web tool[92] (https://string-db.org/) was used to analyze the protein-protein interaction network and the enriched KEGG pathways and Reactome pathways.

### Single-cell trajectory analysis

Mouse normal and fibrotic kidney cells underwent pre-processing and analysis with monocle3[93] using default parameters to generate a single-cell trajectory and dissect cellular decisions, where uninjured VSMCs were considered as the root to perform the pseudo-time analysis.

### Statistical analysis

R (version 4.1.1 and 4.1.3) and GraphPad Prism 8.0.1 were used for the statistical analyses. Differences between two groups were determined using Wilcoxon test or Welch test; P < 0.05 was considered to indicate a significant difference.

## Declarations

### Ethics approval and consent to participate

Not applicable.

### Consent for publication

Not applicable.

### Availability of data and materials

The previously published scRNA-seq data of normal and fatty liver transplantation was deposited at NGDC-GSA under the accession number CRA004061 (https://ngdc.cncb.ac.cn/gsa/browse/CRA004061). Human scRNA-seq datasets of cholangiocarcinoma, bladder cancer, glioblastoma, and ovarian cancer can be accessed through GSE138709, GSE130001, GSE84465, and GSE165897, respectively. The other original data used in this paper can be accessed through the following links: Mouse scRNA-seq dataset of kidney fibrosis (https://zenodo.org/record/4059315), TCGA and CPTAC GBM datasets containing the miRNA and mRNA information (https://www.cbioportal.org/study?id=gbm_tcga_pub2013) and (https://www.cbioportal.org/study?id=gbm_cptac_2021). All other relevant data supporting the key findings of this study are available within the article and its Supplementary Information files or from the corresponding author upon reasonable request. Source codes for the miRTalk R package, the related scripts, and the miRTalkDB database are available at GitHub (https://github.com/multitalk/miRTalk).

### Competing interests

The authors declare that they have no competing interests.

## Funding

This work was supported by the “Pioneer” and “Leading Goose” R&D Program of Zhejiang (2024C03106, X.F.), the National Natural Science Foundation of China (82200725, X.S.; U23A20513, X.F.; 81930016, X.X.), the Key Research and Development Program of China (2021YFA1100500, X.X.), and the Fundamental Research Funds for the Central Universities (226-2024-00001, X.F.)

## Authors’ contributions

X.F., X.X., and X.S. conceived and designed the study. X.S., C.L., and H.Y. developed the miRTalk algorithm. C.L., J.Q., X.Y., and J.L. collected and processed the real-world data. X.S. performed the benchmark test and analyzed the application cases with miRTalk and drew the main figures and supplementary figures. X.S. wrote the manuscript. X.S., X.X., and X.F. revised the manuscript. All the authors read and approved the final manuscript.

## Acknowledgments

The authors thank the High Performance Computing Cluster of Zhejiang University Innovation Center of Yangtze River Delta for their technical support.

## Supplementary Figures

**Fig. S1.**
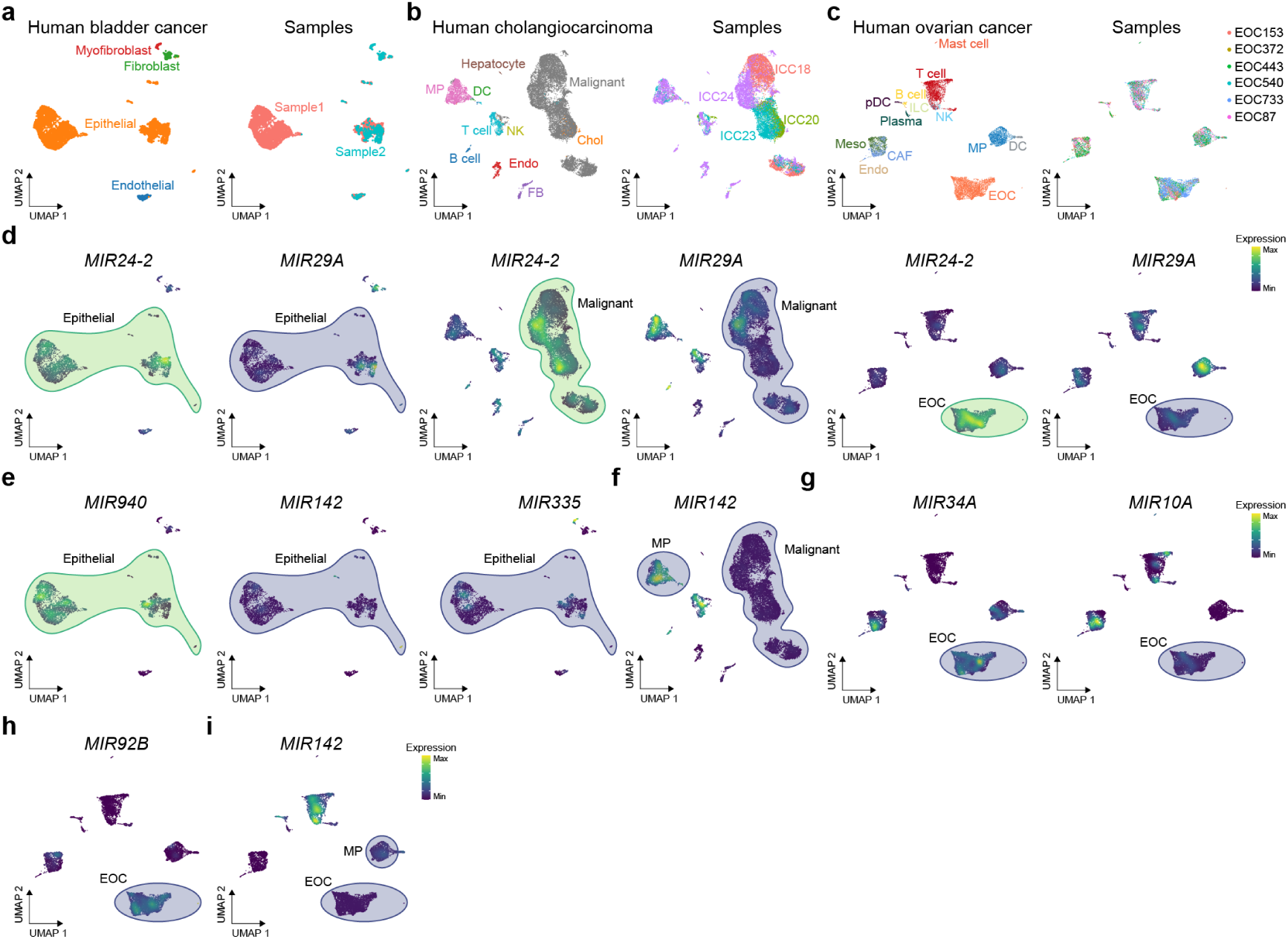
Benchmark on real datasets. **(a)** Human bladder cancer scRNA-seq data including epithelial cells, endothelial cells (Endo), fibroblasts (FB), and myofibroblasts from 2 samples. **(b)** Human cholangiocarcinoma scRNA-seq data including malignant cells, hepatocytes, B cells, T cells, Endo, FB, natural killer (NK) cells, macrophages (MPs), dendritic cells (DCs), and cholangiocytes from 4 samples. **(c)** Human ovarian cancer scRNA-seq data including epithelial ovarian cell (EOC), mast cells, T cells, B cells, plasmacytoid DC (pDC), plasma cells, Endo, mesothelial cells, MPs, DC, innate lymphoid cells (ILCs), cancer-associated fibroblasts (CAFs) from 6 samples. **(d)** Expression of *MIR24-2* and *MIR29* in the human bladder cancer, cholangiocarcinoma, and ovarian cancer. **(e)** Expression of *MIR940, MIR142*, and *MIR335* in the human bladder cancer. (f) Expression of *MIR142* in the human cholangiocarcinoma. (g-i) Expression of *MIR34A, MIR10A*, *MIR92B*, and *MIR412* in the human ovarian cancer.

**Fig. S2.**
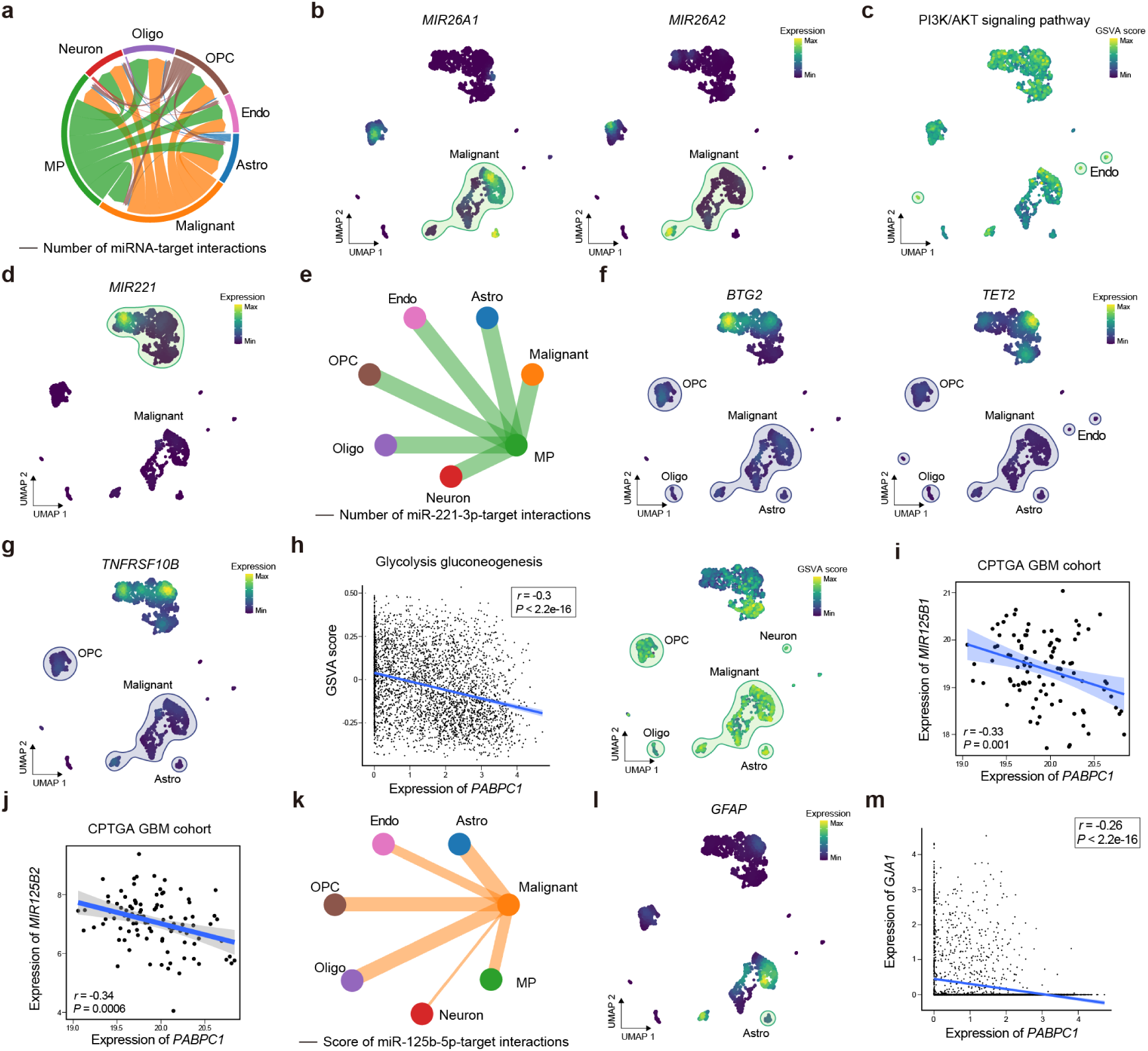
Application of miRTalk in human glioblastoma scRNA-seq data. **(a)** EV-derived miRNA-mediated cell-cell communications inferred by miRTalk, delineating the number of miRNA-target interactions amid pairwise cell types in GBM. **(b)** Expression of *MIR26A1* and *MIR26A2* that encode the miR-26a-5p shown in the UMAP plot. **(c)** GSVA score of the PI3K/AKT signaling pathway. **(d)** Expression of *MIR221* that encode the miR-221 shown in the UMAP plot. **(e)** Number of inferred miR-221-3p-target interactions from MPs to other cell types. **(f-g)** UMAP plot displaying the expression of *BTG2*, *TET2*, and *TNFRSF10B*. **(h)** Inverse correlation between the GSVA score of the glycolysis-gluconeogenesis and the expression of *PABPC1* analyzed with the Pearson’s coefficient. **(i-j)** Verification of the negative association between the expression of *MIR125B1*, *MIR125B2*, and *PABPC1* within the CPTGA GBM cohort. **(k)** Score of inferred miR-125b-5p-target interactions from malignant cells to other cell types. **(l)** UMAP plot displaying the expression of *GFAP*. **(m)** Negative correlation between the expression of *PABPC1* and *GJA1*, assessed utilizing the Pearson’s coefficient.

**Fig. S3.**
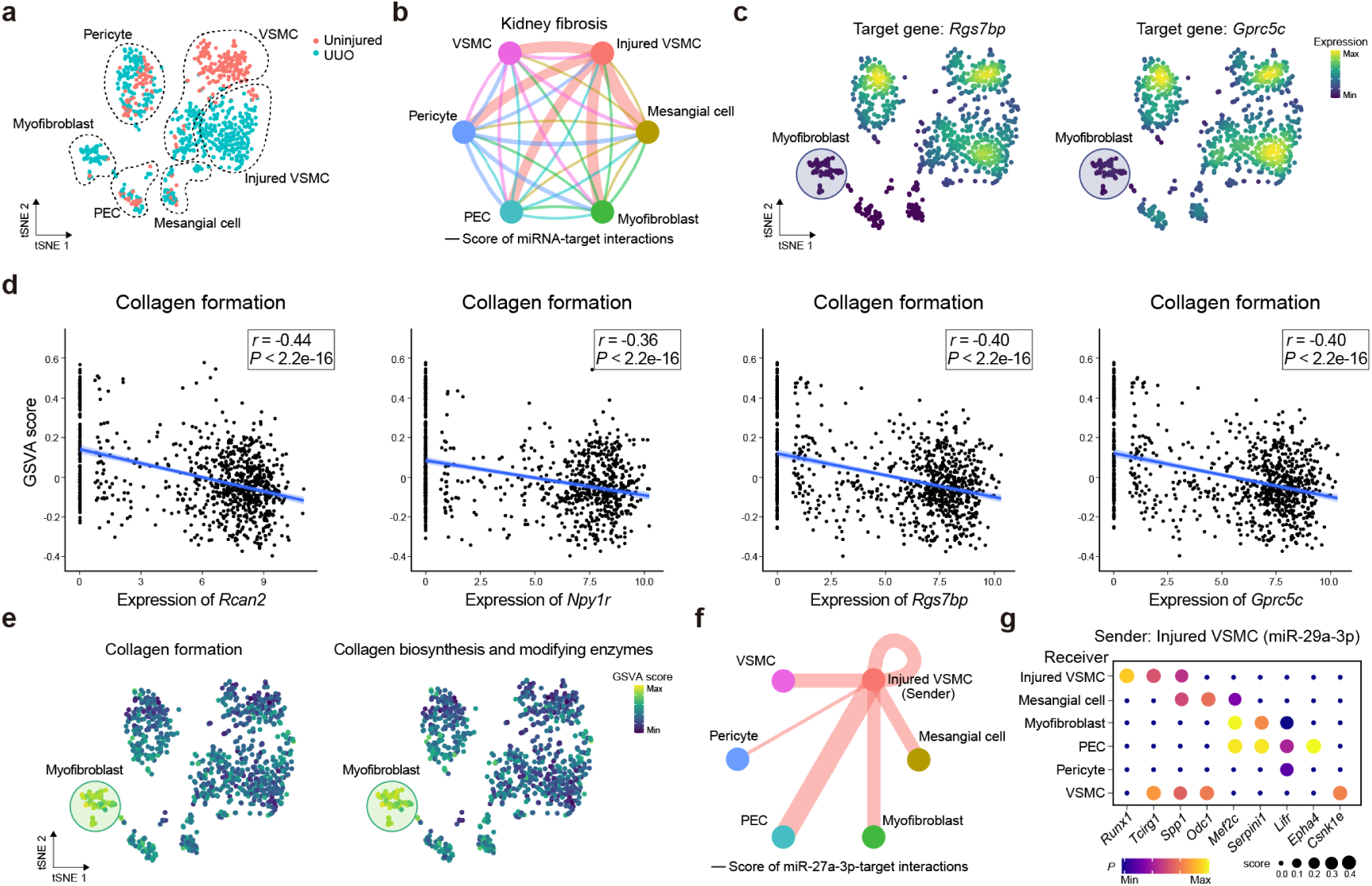
Application of miRTalk in mouse scRNA-seq data of fibrotic kidney. **(a)** UMAP plot displaying the kidney cells at the stage of uninjured and UUO. **(b)** EV-derived miRNA-mediated cell-cell communications inferred by miRTalk, displaying the score of miRNA-target interactions between pairwise cell types in fibrotic kidney. (c) UMAP plot showing the expression of target genes, namely *Rgs7bp* and *Gprc5c*. **(d)** Negative correlation between the GSVA score of collagen formation and the expression of *Rcan2*, *Npy1r*, *Rgs7bp*, and *Gprc5c* analyzed with the Pearson’s coefficient. **(e)** UMAP plot displaying GSVA scores for the collagen formation pathway and the pathway of collagen biosynthesis and modifying enzymes. **(f)** Autocrine and paracrine communications mediated by the injured VSMC-derived miR-27a-3p, where the miRNA-target interaction score was displayed. **(g)** Autocrine and paracrine communication scores and probabilities of inferred miR-27a-3p-target interactions from injured VSMCs to other cell types.

**Fig. S4.**
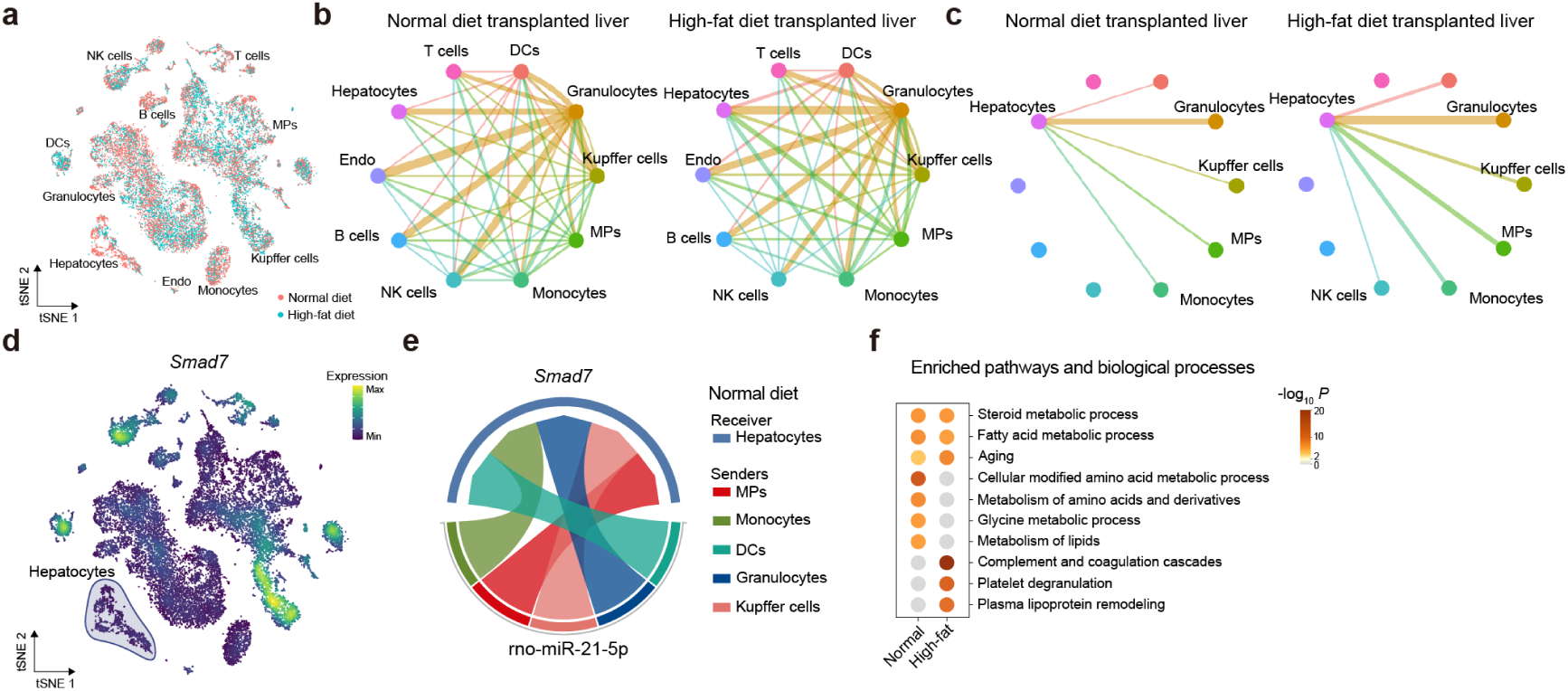
Application of miRTalk in rat scRNA-seq data of normal and high-fat diet liver transplantation. **(a)** UMAP plot displaying the rat liver cells at the condition of normal and high-fat diet. **(b)** EV-derived miRNA-mediated cell-cell communications inferred by miRTalk, displaying the score of miRNA-target interactions between pairwise cell types in normal and high-fat diet transplanted livers. **(c)** EV-derived miRNA-mediated cell-cell communications inferred by miRTalk, displaying the score of miRNA-target interactions from other cells to hepatocytes. (d) Expression of *Smad7* in hepatocytes targeted by miR-21-5p. **(e)** The interaction between miR-21-5p and its target gene *Smad7* that mediates the communication from other cells to hepatocytes in normal livers. **(e)** Enriched pathways and biological processes with the DEGs of hepatocytes between the normal and high-fat diet transplanted livers.

## Notes

https://github.com/multitalk/miRTalk

### Competing Interest Statement

The authors have declared no competing interest.

